# Mutations in tau protein promote aggregation by favoring extended conformations

**DOI:** 10.1101/2023.05.12.540512

**Authors:** Kevin Pounot, Clara Piersson, Andrew Goring, Martin Weik, Songi Han, Yann Fichou

## Abstract

Amyloid aggregation of the intrinsically disordered protein (IDP) tau is involved in several diseases, called tauopathies. Mutations in the gene encoding tau are responsible for a class of inherited tauopathies called frontotemporal dementia and parkinsonism linked to chromosome 17Q (FTDP-17). These mutations are thought to trigger FTDP-17 by favoring the formation of tau amyloid fibrils. This work aims at deciphering the mechanisms through which the diseases-associated single point mutations promote amyloid formation. We combined biochemical characterization and small angle X-ray scattering (SAXS) to study six different FTDP-17 derived mutations. We found that the mutations promote aggregation to different degrees and can modulate tau conformational ensembles, intermolecular interactions and liquid-liquid phase separation propensity. In particular, we found a good correlation between the aggregation lag time of the mutants and their radius of gyration. We show that mutations disfavor intramolecular protein interactions which in turn favor extended conformations and promote amyloid aggregation. This work proposes a new connection between the structural features of tau monomers and their propensity to aggregate, providing a novel assay to evaluate aggregation propensity of tau variants.

## 1. Introduction

Tau pathologies form a class of neurodegenerative diseases in which deleterious deposits enriched in a protein called tau are present in the brain. The tau protein accumulates in these deposits in the form of amyloid filaments, which are highly ordered protein aggregates in which each protein stacks in a cross-beta structure. Strikingly, recent structural work pointed toward a correlation between the conformation of tau within these amyloid aggregates and the associated pathology phenotype ^1^.

Several disease-associated mutations have been identified in the tau proteins, in particular in frontotemporal dementia and parkinsonism (FTDP) linked to chromosome 17 (FTDP-17) (see review by Goedert and Jakes ^2^). Most of the mutations are present in the repeat domains, i.e. in the region that both binds to microtubule and forms the core of amyloid filaments. Accordingly, mutations can exhibit a dual effect on the protein activity : they hinder microtubule binding ^3–5^, and they promote amyloid assembly ^5–8^. The latter findings result from studies conducted in different conditions – inducers, incubation, buffer – and are thus not amenable to a quantitative comparison across mutants. The lack of consistency makes the evaluation of the effect of a specific mutation difficult.

Tau is an intrinsically disordered protein (IDP), meaning that it does not possess a well-defined 3D structure, but rather coexists as many different conformations. In solution state, tau lacks any stable secondary structure elements and is mostly in random coil conformation ^9^. This limits the applicable biophysical methods and has hindered the understanding of the relationship between tau conformations and tau aggregation propensity. The mutation P301L was first shown by NMR to have a small but significant effect on the local conformation with no increased beta-sheet propensity ^10^. Early work suggested that mutations modify the conformation of the flanking regions of aggregation prone regions PHF6 (306-311) and PHF6^*^(275-280) ^6^. More details were provided by Chen *et al*. showing that unshielding the PH6 region can explain mutation-induced aggregation enhancement ^11^. More generally, long-range intramolecular interactions seem to play an important role in aggregation modulation as shown by FRET and cross-linking mass spectrometry ^11–13^. Yet, there is no quantitative work linking the modulation of aggregation propensity by mutations and their structural properties.

Here we studied the effect of six different disease-associated mutations on aggregation propensity and structural features of tau. We used a fragment of tau, referred to as tau187 (residues 255-441 of full length 2N4R, Fig. 1), onto which we engineered the following single point mutations: I260V, G272V, P301L, P301S, Q336R and V337M (Fig. 1). Tau187 contains most of the four repeat domains and the C terminal region, thus encompassing the known amyloid cores, while being more prone to aggregate as compared to full-length 2N4R. Using small-angle X-ray scattering (SAXS) and biochemical characterization methods, we show that a single point mutation modulates tau monomer conformation, intra- and inter-protein interactions and aggregation propensity. We find a good correlation between aggregation lag time and the radius of gyration.

**Figure 1:**
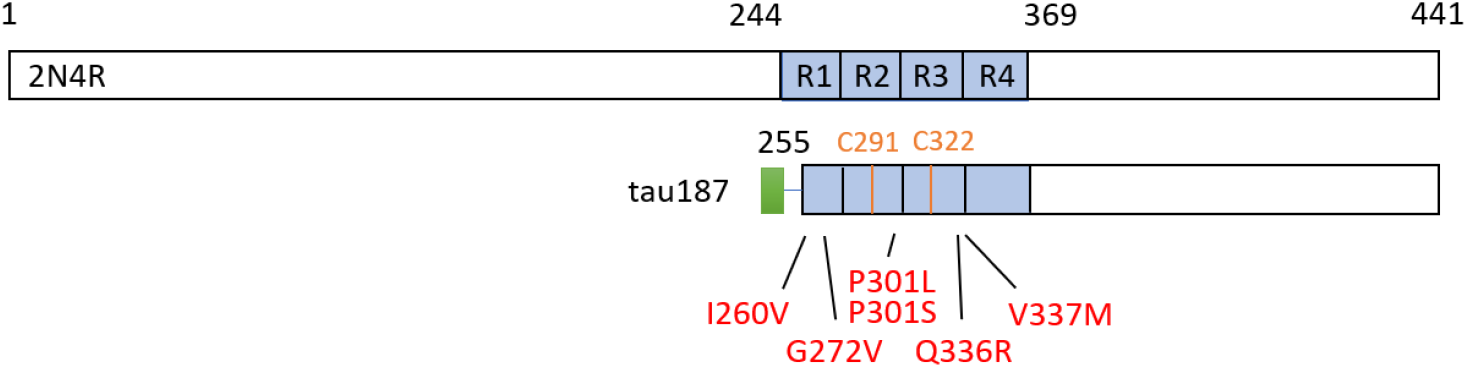
The longest human tau isoform 2N4R contains 441 amino acids. In this work we have used a fragment of 2N4R that starts at residue 255, referred to as tau187, onto which were added single point mutations. Each mutation (red) is located in one of the repeat domains R1 to R4. The green region indicates a poly-histidine tag.

## 2. Results

From the construct tau187, referred to as tau187 WT, we made six mutants that each contained a disease-associated single-point mutation: tau187-I260V, tau187-G27V, tau187-P301L, tau187-P301S, tau187-Q336R, tau187-V337M (Fig. 1).

### 2.1. Single point mutations have an important effect on aggregation kinetic

Amyloid aggregation for all mutants was assessed by Thioflavin T (ThT) fluorescence and transmission electron microscopy (TEM). All mutants were stable over 4 days at 37 °C under shaking (Fig. S1). The addition of a cofactor, the RNA homonucleotide polycytosine (polyC), was used to favor the formation of amyloid fibrils on the experimental timescale.

We assessed the aggregation propensity by recording the ThT fluorescence as a function of time where the RNA was added at time t = 0 h (Fig. 2A). All ThT curves were fitted with a sigmoid function (see Materials and Methods) from which was extracted the aggregation half time (Fig. 2B). The single-point mutations show a drastic effect on aggregation kinetics. All mutations enabled aggregation to occur over 110 h, which was not seen in the WT version. The P301S and P301L mutations lead to the fastest aggregation. Aggregates were observed by TEM for all mutants (Fig. S2).

**Figure 2:**
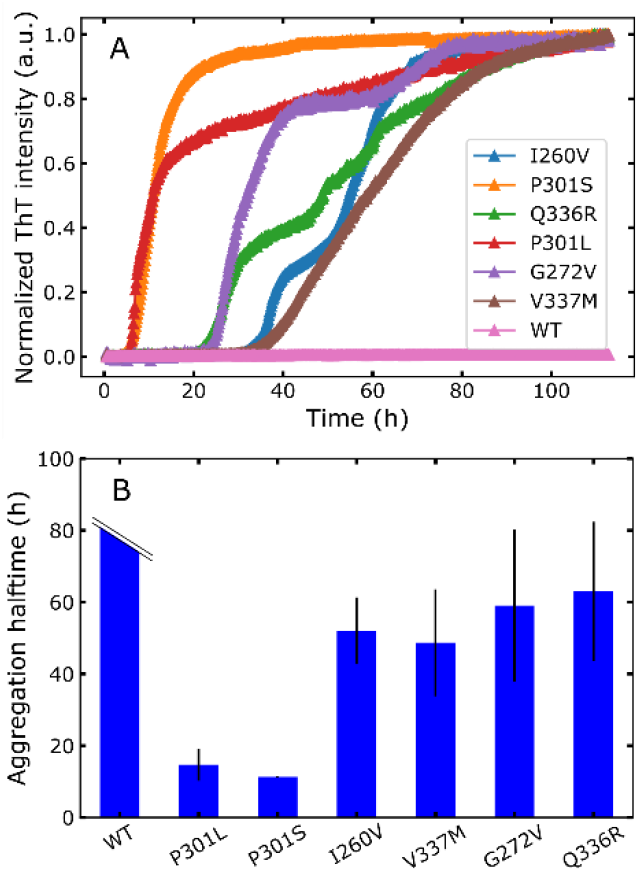
ThT fluorescence as a function of time for different mutants of tau187 incubated with RNA polyC (A). The curves are normalized to their maximum intensity and represent an average of three replicates. Aggregation half time for each mutant (B) extracted from a fit of each ThT curves. Error bars represent the standard deviation over the half times obtained from three independent replicates.

### 2.2. Tau expansion, due to mutations, correlates with aggregation propensity

We then characterized the structural properties of each mutant by small angle X-ray scattering (SAXS). We measured SAXS of all tau187 mutants and extracted their radius of gyration (Fig. 3A). Tau187-WT together with tau187-Q336R exhibit the smallest Rg of 4.1 +/- 0.1 nm. Tau187-P301L and tau187-P301S present the highest Rg of 4.4 +/- 0.06 nm and 4.5 +/- 0.07 nm, respectively. Other mutants exhibit intermediate Rg. These data show that a single-point mutation can significantly change the Rg of the protein, reflecting a change in the conformation ensemble of the different mutants.

**Figure 3:**
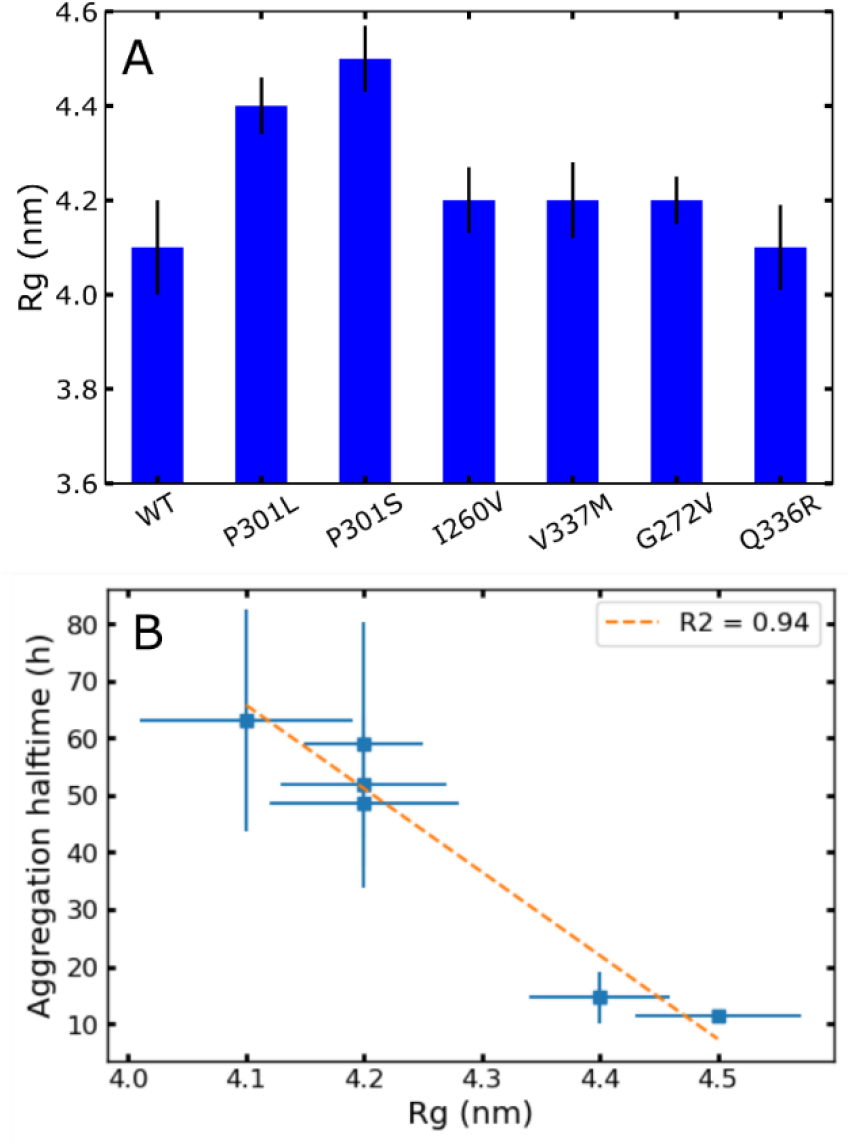
A single point mutation significantly changes tau conformational ensembles, as shown by different radius of gyration Rg (A). The Rg correlates with aggregation half time (B). Error bars on Rg originates from the covariance matrix generated by the fitting procedure. Tau187-WT does not aggregate on the observed timescale and thus has no aggregation half time. Hence it is not plotted here in panel B.

We further investigated whether the aggregation propensity could be linked to structural features of the different mutants. We plotted the aggregation half time as a function of the radius of gyration Rg (Fig. 3B). Strikingly, we found an excellent correlation (R^2^ =0.94) for the two parameters, demonstrating that the more extended a mutant is, the more aggregation prone it is.

### 2.3. Interactions between tau molecules are repulsive and modulated by single point mutation

Aggregation involves the assembly of many molecules together. Thus, we investigated the nature of interactions between tau molecules in solution. Light scattering from a particle solution intrinsically contains information on the interactions between the particles. Here, we extracted from SAXS experiments the second virial coefficient A_2_, a parameter that reflects interparticle interactions ^14^. The scattering intensities extrapolated at scattering angle q=0 were plotted against protein concentration (Fig. S3) and fitted with equation 1 (see Materials & Methods). Positive values of A_2_ originate from repulsive interactions (the higher A_2_ is, the more repulsion exists) and negative A_2_ originate from attractive interactions (the more negative A_2_ is, the more attraction exists). Figure 4A presents A_2_ for each mutant. A_2_ is significantly different for all mutants, revealing that a single point mutation is sufficient to significantly modulate intermolecular protein interactions. In addition, A_2_ are positive for all mutants but Q336R, pointing to the fact that interactions are overall repulsive between tau molecules. This is in good agreement with the observation that recombinant tau is extremely stable in solution and did not aggregate in the absence of RNA (Fig. S1). Moreover, we evaluated if the interaction between tau monomers is linked to aggregation propensity triggered by RNA. We found no correlation between A_2_ and aggregation halftime (Fig. 4B), confirming that aggregation induced by RNA is not directly related to intermolecular protein interactions in solution.

**Figure 4:**
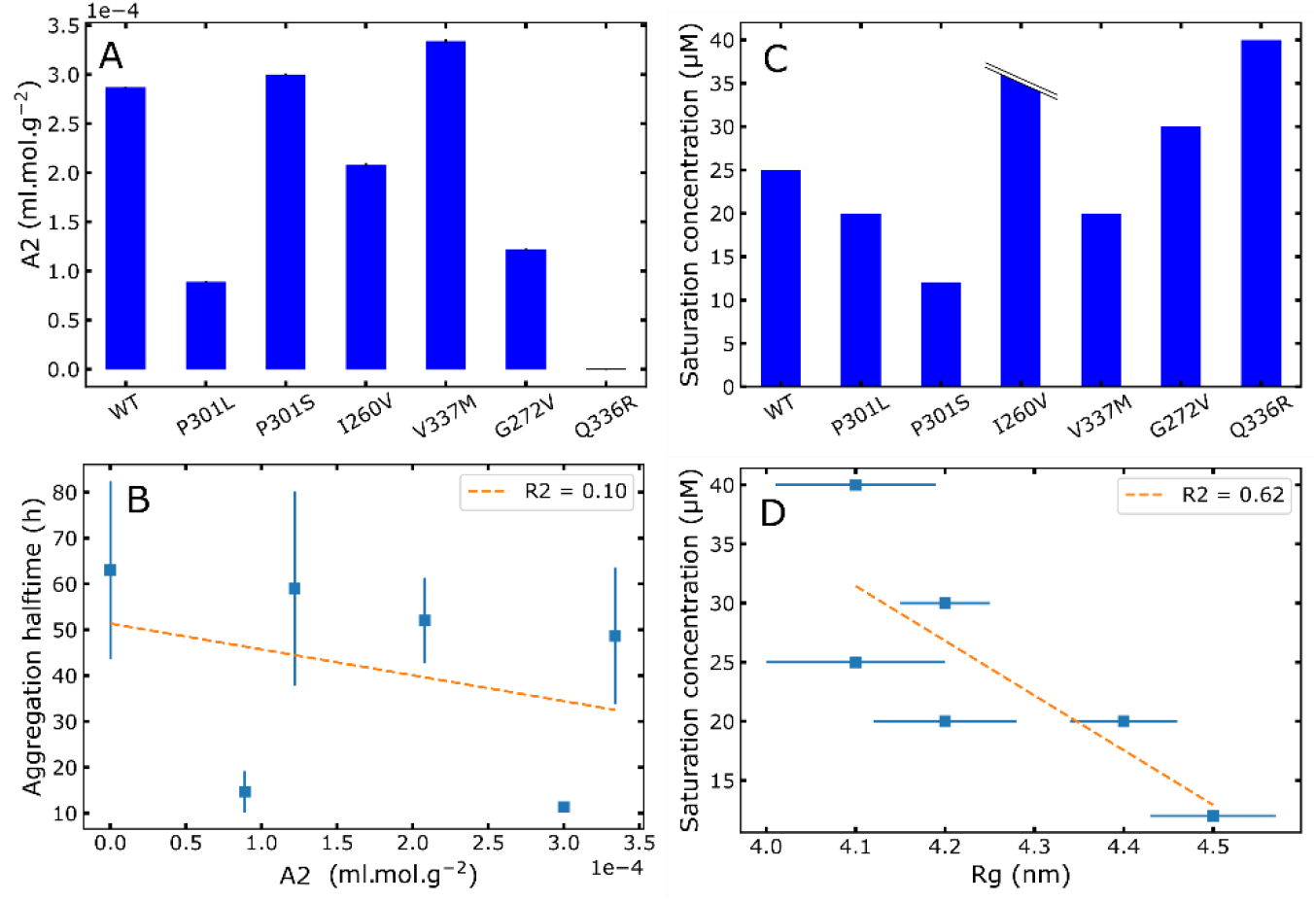
Inter-protein interactions modulated by mutations. (A) The second virial coefficient A_2_ shows overall repulsive interactions. Error bars are from the fitting procedure. (B) Aggregation half time is not linked A_2_. (C) LLPS saturation concentration C_sat_ for different mutants (I260V exceeded the tested maximum concentration of 40 uM). C_sat_ is defined as the lowest concentration giving a significant absorption over 3 independent replicates (p-value < 0.05; see methods) (D) LLPS saturation concentration correlates with Rg. For correlation plots (B) and (D), WT and I260V data points are not shown because their aggregation half time and saturation concentration, respectively, are not defined.

Next, we evaluated another parameter reflecting inter-molecule interactions that is the capacity to participate in liquid-liquid phase separation (LLPS). LLPS is a physical process where protein molecules can spontaneously form a high concentration phase relying on a network of interaction between the different molecules ^15^. Tau was previously shown to form LLPS under high salt conditions, where hydrophobic interactions are enhanced ^16^. In a buffer containing 3 M NaCl, we measured the saturation concentration (C_sat_), which is the minimum protein concentration at which the formation of LLPS is observed (Fig. S4). Figure 4C shows the C_sat_ for all mutants. All mutants but I260V exhibited LLPS in the explored range of 0-40 μM. We found that C_sat_ is modulated by the single-point mutations. Tau187-P301L, tau187-P301S and tau187-V337M are more prone than tau187-WT to form LLPS, suggesting that these mutations favor hydrophobic intermolecular protein interactions. In contrast, tau187-I260V (which did not form LLPS up to 40 μM), tau187-G272V and tau187-Q336R are less prone than tau187-WT to form LLPS. Furthermore, we found a good correlation between the propensity to form LLPS, as shown by the saturation concentration, and Rg (Fig 4D). This correlation indicate that more extended conformations facilitate hydrophobic interactions, which are responsible for high-salt LLPS.

### 2.4. Extended conformations originate from enhanced protein-water interactions

Then we further investigated the origins of the variation of tau conformations in the different mutants. To do so, we evaluated the contributions of intramolecular protein-protein interactions and protein-solvent interactions, we treated the SAXS data with a previously developed approach aiming at assessing the hydration quality of disordered proteins ^17,18^. From polymer theory, the radius of gyration typically follows the relation Rg ∼ N^ν^, where N is the length of the polymer and *v* the Flory exponent. Riback et al. used simulations on model polymers to obtain the relation between Rg and ν for various solvent conditions ^17^. The polymer molecular form factor and the Flory exponent ν were extracted from the SAXS data using the web server made available by Riback et al. (Fig. S5). The ν parameter reflects the protein solvation quality for disordered polymers: ν is greater or smaller than 0.5 for favored or disfavored protein-solvent interactions, respectively. Figure 5A shows *ν* for the different mutants. Note that tau187-WT data produced a poor-quality fit and therefore its Flory exponent was considered not reliable and was not analyzed (see Fig. S6). We observe in Fig. 5A that the quality of hydration is significantly modulated by the single point mutations. P301L/S show the highest ν, which suggests from polymer theory that these mutations favor protein-solvent interactions over protein-protein interactions. The value of ν is furthermore positively correlated with Rg (Fig. 5B). This correlation shows that weaker intramolecular protein interactions (i. e. higher Flory exponent) promote extended conformations, and in turn leads to increased aggregation propensity (fig. 5C).

**Figure 5:**
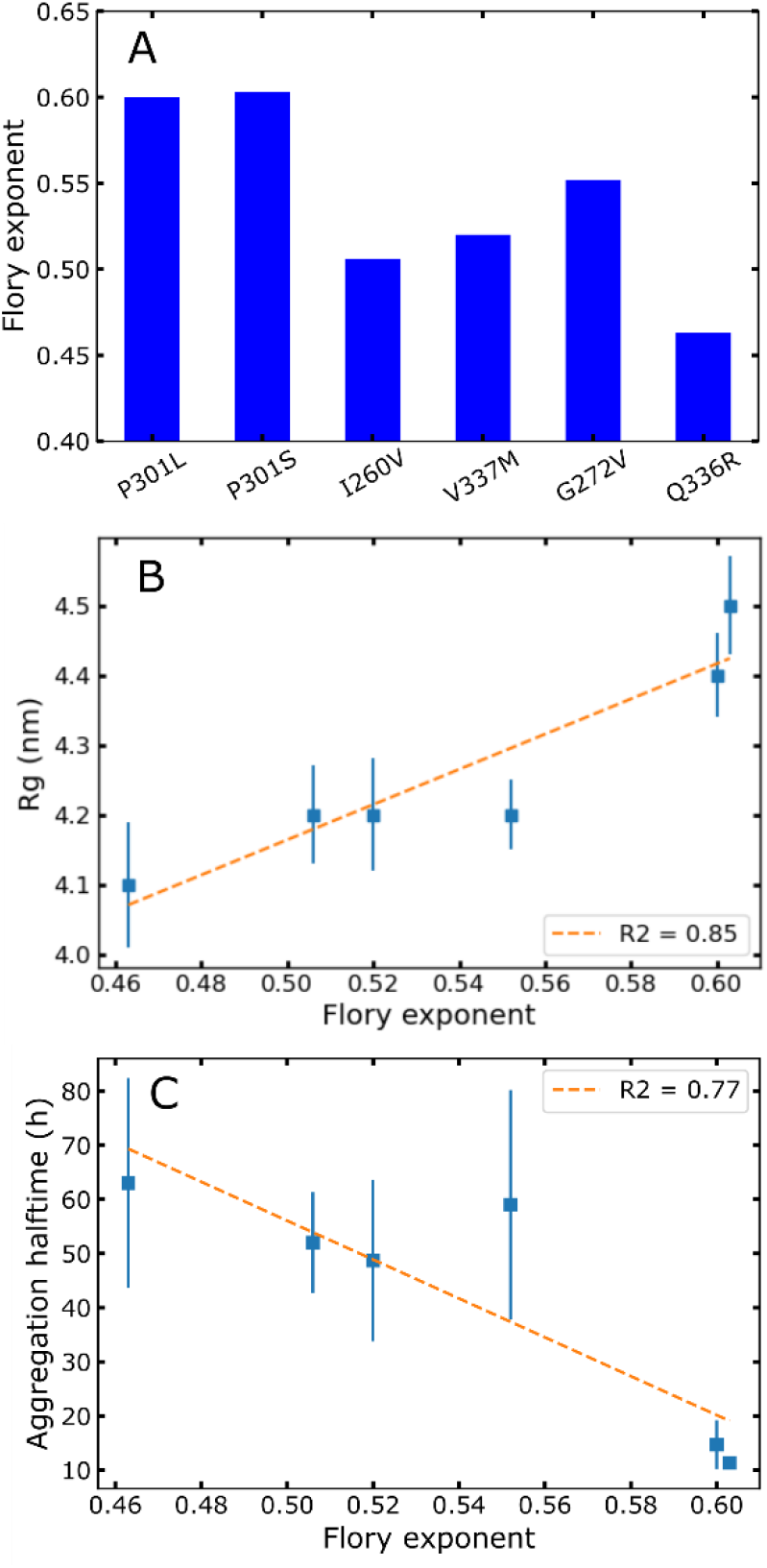
(A) Flory exponent ν of tau187 mutants reveal that single point mutations modulate tau-solvent interactions. (B) Rg is positively correlated with ν and (C) aggregation half time is negatively correlated with ν.

## 3. Discussion

We analyzed the aggregation propensity and structural features of different disease-associated mutants of the tau187 protein fragment. We showed that single point mutations significantly change the aggregation propensity, as observed by different aggregation half time in the presence of RNA as an inducer. We further found that single point mutations change tau conformations of as well as the interactions between monomers. Strikingly we found that the measured Rg correlates with aggregation half time.

Early studies suggested that structural effects of disease-associated mutations on tau monomers were small or nonexistent ^6,7,10^. NMR showed subtle local structural changes that were hard to interpret in terms of conformational changes ^10^. A study of the P301L mutation by Chen et al. provided a more comprehensive view of the structural effect of the mutation. They showed that the P301L mutation induces a local opening of the protein, thereby exposing a specific amyloidogenic region of tau. In agreement with this report, here we show that more generally, an extension of the protein, as probed by its Rg, directly correlates with the aggregation half time of a given mutant (Fig. 6). This finding is in good agreement with the model where extended conformers are overall favorable for intermolecular contacts of the amyloidogenic region such as PHF6 and PHF6^*^.

**Figure 6:**
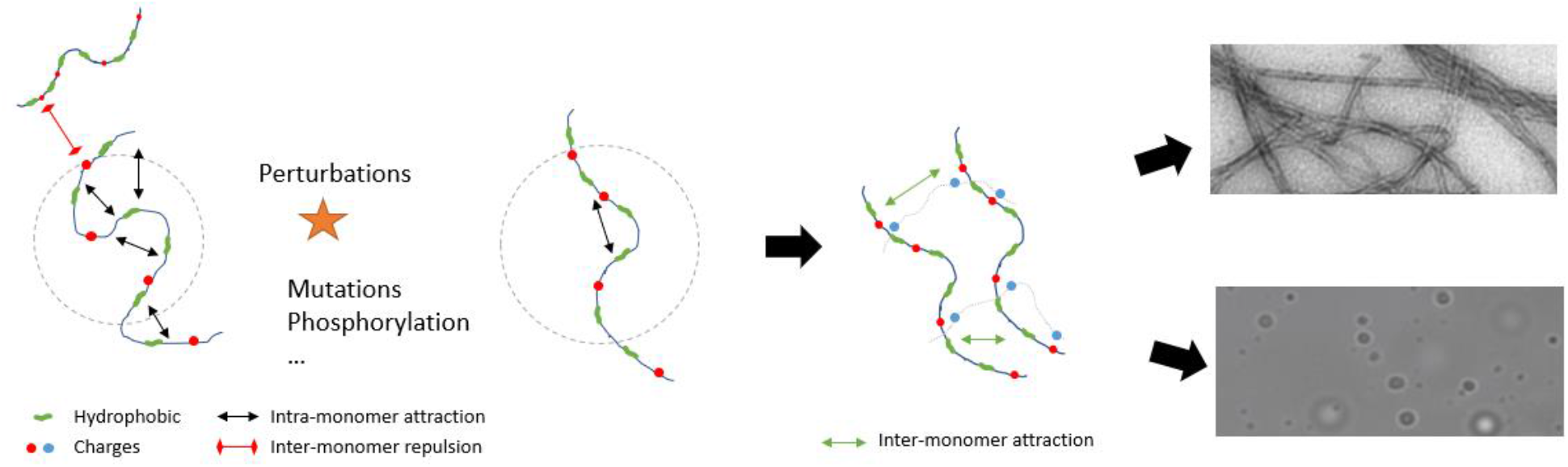
Aggregation model based on the presented data. In solution, the tau proteins exhibit intermolecular repulsions and does not readily aggregate. Upon modifications, such as mutations, protective intramolecular interactions are released, leading to an increased radius of gyration. These extended conformations are more prone to intermolecular hydrophobic interactions, which in turn drive the formation of amyloid aggregates and LLPS.

The analysis of the full SAXS curve using a molecular form factor for disordered polymer ^17^ (Fig. 5 and S5) allowed us to extract the Flory exponent, reflecting the quality of the protein solvation. We show that mutations modulate this parameter. Specifically, some mutants (most drastically P301L/S) exhibit enhanced protein-solvent interactions over intramolecular protein interactions. Although one might intuitively think that favoring solvation quality would stabilize the protein, here we highlight a different mechanism (Fig. 6). Rather, P301L/S mutations disrupt protective intramolecular interactions observed before ^11^, thereby leading to increased radius of gyrations and higher aggregation propensities. This mechanism is furthermore consistent with the observation that an increased Rg correlates with high-salt LLPS propensity (Fig 4D) which relies on hydrophobic interprotein contacts ^16^. Notably, a similar mechanism has been highlighted for another amyloid-forming IDP, the α-synuclein ^19^. More generally, this work demonstrates that tau aggregation propensity is encoded, at least partially, in the monomer structure, in agreement with reports on other IDPs such as α-synuclein ^19^ and Huntingtin ^20^.

Analysis of A_2_ showed that overall the tau monomers exhibit repulsive interactions (Fig. 4A). This observation might explain why recombinant tau hardly aggregates *in vitro* and often requires the use of inducers such as heparin or RNA. The observation that the aggregation half time induced by RNA does not correlate with A2 shows that RNA completely rewrites this repulsion, likely by compensating the numerous charges present in the tau protein. One can speculate that cofactor free aggregation seen for smaller fragments ^11,21^ would be more related to this A2 parameter. RNA acts as a “mild” cofactor ^22^ so that the protein still needs to overcome a significant energy barrier to form amyloid, leading to a significant lag time (Fig. 2A). Herein this work we show that increasing the population of aggregation-prone conformers, characterized by lower intramolecular affinity and increased Rg, reduces this lag time and therefore the energy barrier to form ThT active species. The impact of mutations is not expected to be similar for other conditions or cofactors, such as heparin, that completely suppress this lag time. Indeed, we verified that there is no significant lag time for the different mutants incubated with heparin and that aggregation half time does not correlate with Rg (Fig S7). In this fast kinetics without lag phase, the aggregation half time is dominated by the growth rate, which we conjecture to be very dependent on the detailed properties of the tau-cofactor interactions. This view is in line with a recent report showing that the effect of mutations on aggregation is inducer dependent ^23^.

A direct correlation between aggregation half time and Rg lays the ground for a rapid and convenient way to evaluate tau aggregation propensity. From SAXS measurements, one can obtain Rg, A2 and ν of different tau variants, e.g. carrying different post-translation modifications or mutations, to obtain an idea of the aggregation propensity of the variant. This is particularly useful as many different combinations of modifications such as phosphorylation are possible and have been shown to have non-trivial effect on aggregation propensity ^24^.

## 4. Material and methods

### 4.2. Protein expression and purification

Tau187, a truncated version of 2N4R (residues 255-441) was engineered with a poly-histidine tag at the N-terminal end. Mutants of tau187 were prepared using site-directed mutagenesis.

The expression and purification of tau187 variants have been previously reported ^25,26^. Genes were transformed into E. Coli BL21(DE3) that grew at 37 °C, 200 rpm, with addition of 10 μg/mL kanamycin, until reaching optical density (600nm) of 0.6. Expression was induced by incubation with 1 mM isopropyl- ß-D-thiogalactoside for 2–3 h. Cells were harvested with centrifugation at 5000 g for 20 min. Cell pellets were resuspended in lysis buffer (Tris-HCl, pH = 7.4, 100 mM NaCl, 0.5 mM DTT, 0.1 mM EDTA) added with 1 Pierce protease inhibitor tablet (Thermo Scientific, A32965), 1 mM PMSF, 2 mg/mL lysozyme, 20 μg/mL DNase and 10 mM MgCl_2_ (10 mM), and incubated on ice for 30 min. Samples were then frozen and thawed for 3 times using liquid nitrogen, then centrifuged at 10,000 rpm for 10 min. 1 mM PMSF was added again and samples were heated at 65 °C for 12 min and cooled on ice for 20 min. Cooled samples were then centrifuged at 10,000 rpm for 10 min to remove the precipitant. The resulting supernatant was loaded onto a column pre-packed with 5ml Ni-NTA resins (cytivia HisTrap HP) using an Akta pure system. The column was washed with 25 mL of buffer A (20 mM sodium phosphate, pH = 7.0, 500 mM NaCl, 10 mM imidazole, 100 μM EDTA), 25 mL of buffer B (20 mM sodium phosphate, pH = 7.0, 1 M NaCl, 20 mM imidazole, 0.5 mM DTT, 100 μM EDTA) and 25 ml of buffer A. Protein was eluted with 0-100% gradient over 50 ml of buffer C (20 mM sodium phosphate, pH = 7.0, 0.5 mM DTT, 100 mM NaCl, 300 mM imidazole). Eluents were analyzed by SDS-PAGE to collect the pure fractions. Proteins were then buffer exchanged into working buffer of 20 mM HEPES, 100mM NaCl, pH 7.0.

### 4.2. SAXS experiments

The SAXS experiments were conducted on the BM29 beamline at the European synchrotron radiation facility (ESRF) ^27,28^. Following two different procedures.

The first experiment was performed in batch, by loading 50 μL of sample in the dedicated BM29 sample changer after centrifugation at 10,000 rpm for 10 minutes. For each sample measured, a series of 20 frames was acquired with an integration time of 1 s for each frame. The azimuthal integration of the images was done automatically from the beamline control software and the 1D scattering curves were subsequently used for data analysis. These datasets were used to extract the radius of gyration and the Flory exponent.

In a second experiment, a size-exclusion chromatography (SEC) S200 column of 3 mL volume was used to separate possible contaminants and aggregates from the monomers. The buffer used was 20mM ammonium acetate, 100mM NaCl and 5mM TCEP. The samples were spin downed at 10,000 rpm for 10 minutes before the injection on the column. The flow rate was set to 0.3 mL/min and the injected volume was 50 μL. The images were acquired on a Pilatus 2M detector at a distance of 2.869 m from the sample. Each image is obtained from a 0.5 s exposure of the sample to the X-ray beam. These data were used to extract A2 parameter using the online UV-VIS spectrophotometer to extract protein concentration and directly obtain I(c,0).

### 4.3. SAXS data treatment and analysis

The radius of gyration (Rg) was obtained from the linearized batch SAXS data, ln(I(q)) vs. q^2^, fitted using the following relation 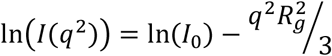. To obtain a reliable value of the Rg, the fit is performed on multiple subdivisions of the fitting region. The subdivisions size ranges from 6 points to the full q-region used for the fit. The obtained q-values are then plotted on a frequency histogram and the final Rg is obtained by taking the weighted-average of the histogram. The quality of the Rg can be assessed by inspection of the histogram, where a proper linear Guinier region should give a narrow distribution of Rg around the mean value. The obtained Rg values were compared with values obtained from other software (ATSAS and Riback and Sosnick’s web server^17,18^) and showed good agreement.

The Flory exponent was obtained by fitting the frames using a χ^2^ -type distance from the average of all frames. All the frames that deviated from 2σ, σ being the standard deviation of χ^2^ distances, were eliminated. The buffer subtraction was then performed and the subtracted data were used as input for the Riback and Sosnick’s web server ^17,18^. The SEC-SAXS data contain the UV measurement as the function of time along with the X-ray scattering images. The frames that pertain to the UV peak were manually selected, as well as the frames that contain only the buffer. The buffer frames are chosen such that they are positioned in time just before the sample frames, which gives the best buffer subtraction (Fig. S8). The SEC-SAXS data were used to compute the second virial coefficient A_2_ according to the following:

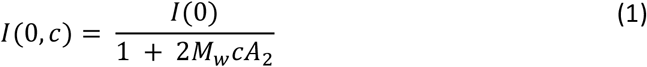

where I(0, c) is the extrapolated SAXS intensity at q = 0 Å^-1^ (obtained from a guinier fit at low q) and at protein concentration c, and M_w_ = 20,570 Da is the protein molecular weight. The concentration was obtained from the UV signal recorded after the SEC by dividing the absorbance at 280 nm by the protein molar extinction coefficient ε = 2,800 M^-1^ cm^-1^. The equation 1 was fitted to the data using the Python Scipy’s *curve_fit* routine.

Scripts for SAXS data analysis are deposited and available on Github (DOI 10.5281/zenodo.7893438).

### 4.4. LLPS experiments

Different concentrations (0-40 μM) of Tau187 mutants were incubated with 3 M NaCl in a 384-well low-volume microplate. Total volume was 30 μl in each well. Each condition was prepared independently in three different wells. Standard deviations over the three wells are presented as error bars. Absorbance at 500 nm was measured in a BMG fluoroStar Omega after 10min of incubation. LLPS was not detected in the range 0-12 μM protein so the absorbances measured at 0, 4, 8 and 12 μM protein were used to define an absorption baseline. A t-test was performed between the absorption of these 4 concentrations and each of the upper concentration. The lowest protein concentration giving p < 0.05 was reported as the saturation concentration. A representative sample of the raw data are shown in Fig. S4 for tau187-WT.

### 4.5. ThT experiments and data fitting

The tau protein was incubated at 20 μM in 384-well low-volume microplate with 20 μM ThT. The RNA polyC (Sigma P4903) was added at 200 μM and the heparin (Sigma H6279) was added at a concentration of 5 μM. The working volume was 20 ul. The fluorescence was bottom read in a BMG fluoroStar Omega with an excitation and emission wavelength of 440 and 480 nm, respectively. Each condition was prepared independently in three different wells.

ThT kinetic curves were fitted using the curve_fit python function with the following equation:

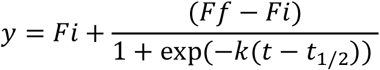

Where Fi and Ff represent the initial and final fluorescent intensities respectively, k represents the growth rate and t_1/2_ represent the aggregation half time. Each well was fitted independently. The presented error bars on the output fit parameters represent the standard deviation over replicates of the same condition. For visualization in Fig. 2A and Fig. S7A, kinetic curves represent the average over 3 replicates and are normalized between 0 and 1.

## Acknowledgements

The author would like to thank Dr. Loquet to provide access to his lab to prepare sample and perform experiments. The authors would like to thank Dr. Riback for discussion about his approach and the webserver. YF would like to thank the European Research Council (Grant 101040138) and the Federation of European Biochemical societies (FEBS) for their financial support.

## References

1. Shi, Y. et al. Structure-based classification of tauopathies. Nature 598, 359–363 (2021).

2. Goedert, M. & Jakes, R. Mutations causing neurodegenerative tauopathies. Biochim. Biophys. Acta 1739, 240–250 (2005).

3. Hong, M. et al. Mutation-Specific Functional Impairments in Distinct Tau Isoforms of Hereditary FTDP-17. Science 282, 1914–1917 (1998).

4. Hasegawa, M., Smith, M. J. & Goedert, M. Tau proteins with FTDP-17 mutations have a reduced ability to promote microtubule assembly. FEBS Lett. 437, 207–210 (1998).

5. Grover, A. et al. A novel tau mutation in exon 9 (1260V) causes a four-repeat tauopathy. Exp. Neurol. 184, 131–140 (2003).

6. von Bergen, M. et al. Mutations of tau protein in frontotemporal dementia promote aggregation of paired helical filaments by enhancing local beta-structure. J. Biol. Chem. 276, 48165–48174 (2001).

7. Goedert, M., Jakes, R. & Crowther, R. A. Effects of frontotemporal dementia FTDP-17 mutations on heparin-induced assembly of tau filaments. FEBS Lett. 450, 306–311 (1999).

8. Nacharaju, P. et al. Accelerated filament formation from tau protein with specific FTDP-17 missense mutations. FEBS Lett. 447, 195–199 (1999).

9. Mukrasch, M. D. et al. Structural Polymorphism of 441-Residue Tau at Single Residue Resolution. PLOS Biol. 7, e1000034 (2009).

10. Fischer, D. et al. Structural and Microtubule Binding Properties of Tau Mutants of Frontotemporal Dementias. Biochemistry 46, 2574–2582 (2007).

11. Chen, D. et al. Tau local structure shields an amyloid-forming motif and controls aggregation propensity. Nat. Commun. 10, 2493 (2019).

12. Jeganathan, S., von Bergen, M., Brutlach, H., Steinhoff, H.-J. & Mandelkow, E. Global Hairpin Folding of Tau in Solution. Biochemistry 45, 2283–2293 (2006).

13. Mirbaha, H. et al. Inert and seed-competent tau monomers suggest structural origins of aggregation. eLife 7, e36584 (2018).

14. Quigley, A. & Williams, D. R. The second virial coefficient as a predictor of protein aggregation propensity: A self-interaction chromatography study. Eur. J. Pharm. Biopharm. Off. J. Arbeitsgemeinschaft Pharm. Verfahrenstechnik EV 96, 282–290 (2015).

15. Alberti, S. & Hyman, A. A. Biomolecular condensates at the nexus of cellular stress, protein aggregation disease and ageing. Nat. Rev. Mol. Cell Biol. 22, 196–213 (2021).

16. Lin, Y. et al. Liquid-Liquid Phase Separation of Tau Driven by Hydrophobic Interaction Facilitates Fibrillization of Tau. J. Mol. Biol. 433, 166731 (2021).

17. Riback, J. A. et al. Innovative scattering analysis shows that hydrophobic disordered proteins are expanded in water. Science 358, 238–241 (2017).

18. Riback, J. A. et al. Commonly used FRET fluorophores promote collapse of an otherwise disordered protein. Proc. Natl. Acad. Sci. 116, 8889–8894 (2019).

19. Stephens, A. D. et al. Extent of N-terminus exposure of monomeric alpha-synuclein determines its aggregation propensity. Nat. Commun. 11, 2820 (2020).

20. Elena-Real, C. A. et al. The structure of pathogenic huntingtin exon 1 defines the bases of its aggregation propensity. Nat. Struct. Mol. Biol. 30, 309–320 (2023).

21. Lövestam, S. et al. Assembly of recombinant tau into filaments identical to those of Alzheimer’s disease and chronic traumatic encephalopathy. eLife 11, e76494 (2022).

22. Fichou, Y. et al. Tau-Cofactor Complexes as Building Blocks of Tau Fibrils. Front. Neurosci. 13, (2019).

23. Ingham, D. J., Hillyer, K. M., McGuire, M. J. & Gamblin, T. C. In vitro Tau Aggregation Inducer Molecules Influence the Effects of MAPT Mutations on Aggregation Dynamics. Biochemistry 61, 1243–1259 (2022).

24. Despres, C. et al. Identification of the Tau phosphorylation pattern that drives its aggregation. Proc. Natl. Acad. Sci. U. S. A. 114, 9080–9085 (2017).

25. Pavlova, A. et al. Protein structural and surface water rearrangement constitute major events in the earliest aggregation stages of tau. Proc. Natl. Acad. Sci. 113, E127–E136 (2016).

26. Fichou, Y., Vigers, M., Goring, A. K., Eschmann, N. A. & Han, S. Heparin-induced tau filaments are structurally heterogeneous and differ from Alzheimer’s disease filaments. Chem. Commun. 54, 4573–4576 (2018).

27. Borowski, M., Bowron, D. T., De Panfilis, S. & IUCr. High-energy X-ray absorption spectroscopy at ESRF BM29. Journal of Synchrotron Radiation vol. 6 179–181 https://scripts.iucr.org/cgi-bin/paper?S0909049599001867 (1999).

28. Pernot, P. et al. Upgraded ESRF BM29 beamline for SAXS on macromolecules in solution. J. Synchrotron Radiat. 20, 660–664 (2013).

