## Supplemental figures for "Mutations in tau protein promote aggregation by favoring extended conformations"

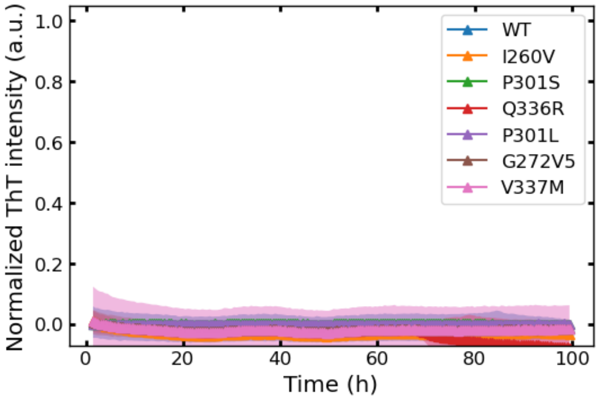


Figure S1 : None of the mutants showed aggregation after 4 days of incubation at 37 °C.

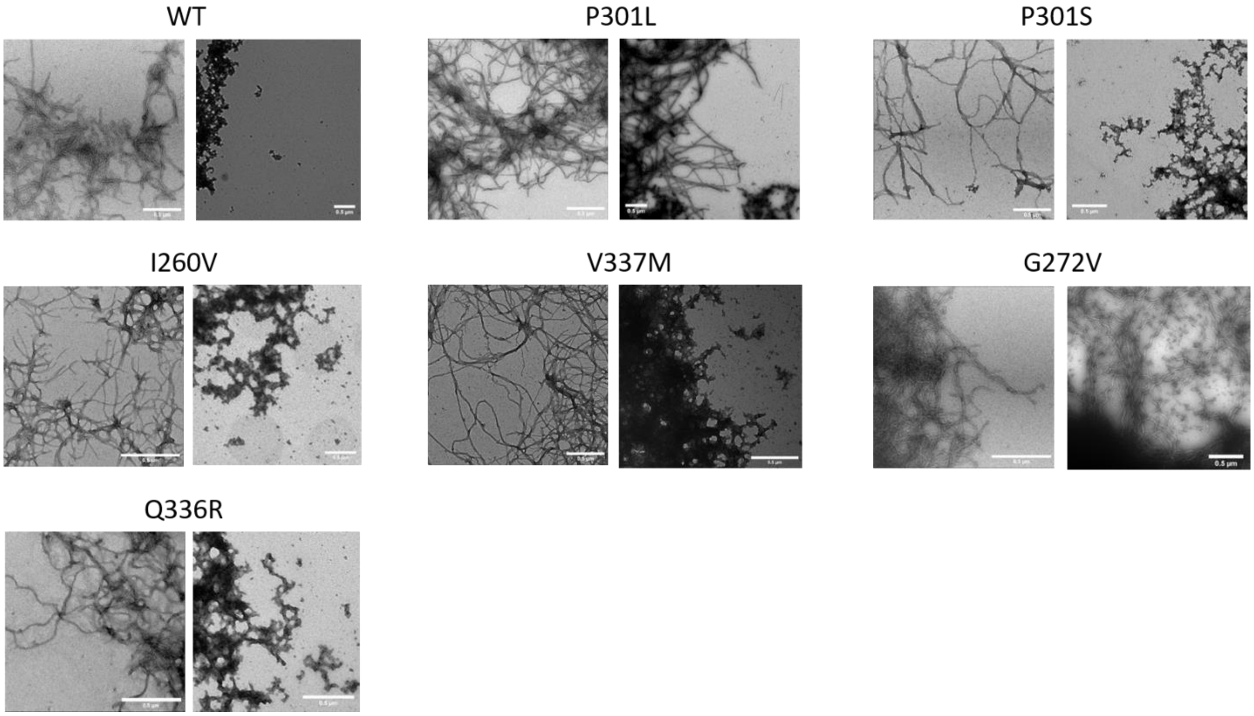


Figure S2 : TEM images for tau187 (20 µM) mutants incubated with heparin (5 µM) for 24h (left) or polyC (200 µg/ml) for 100h (right). In general, poly tau187 incubated with polyC resulted in a mixture of amorphous and filamentous aggregates. Scale bars are 0.5 µm.

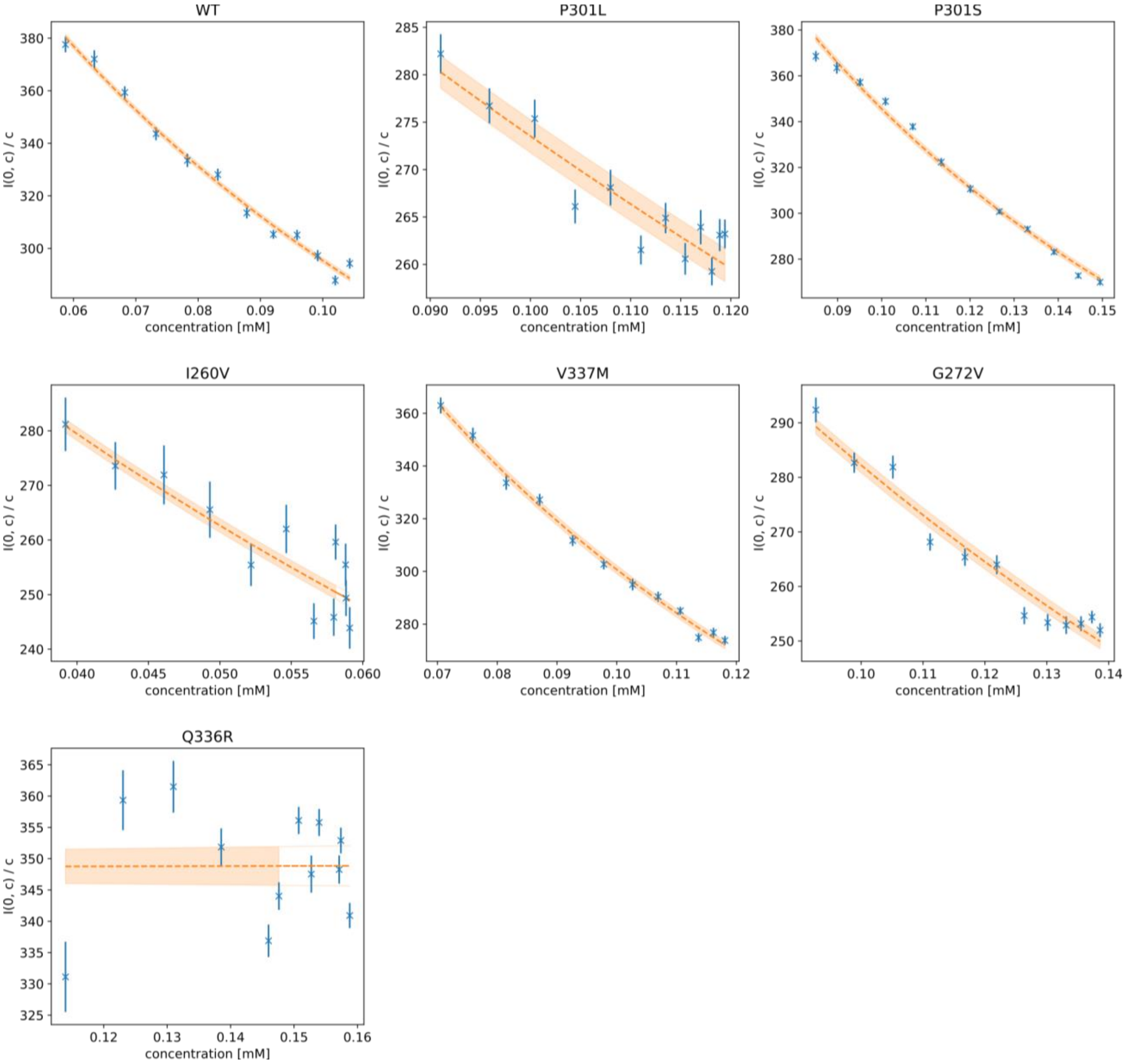


Figure S3 : Fits of I(0) Vs concentration to extract $A_2$. The SEC-SAXS data were treated and analyzed as described in Methods. On each panel, the corresponding mutant is given on top and the extrapolated signal I(0, c) / c (blue crosses with error bar) is plotted as the function of concentration c. The fit of equation 1 is plotted (orange dashed curve) with a confidence interval. The orange area around the fitted curve is the uncertainty on the model computed using:

$$\sigma_f^2 = \left(\frac{\partial f}{\partial I0}\right)^2 \sigma_{I0}^2 + \left(\frac{\partial f}{\partial A_2}\right)^2 \sigma_{A_2}^2 + 2\frac{\partial f}{\partial I0}\frac{\partial f}{\partial A_2} \quad Cov(I0, A_2) \qquad (2)$$

where f is the fitting function from equation 1, $\sigma_{I0}^2$ and $\sigma_{A_2}^2$ are the estimated errors on the fitted parameters for I0 and $A_2$, respectively and Cov(I0, $A_2$) is the covariance function.

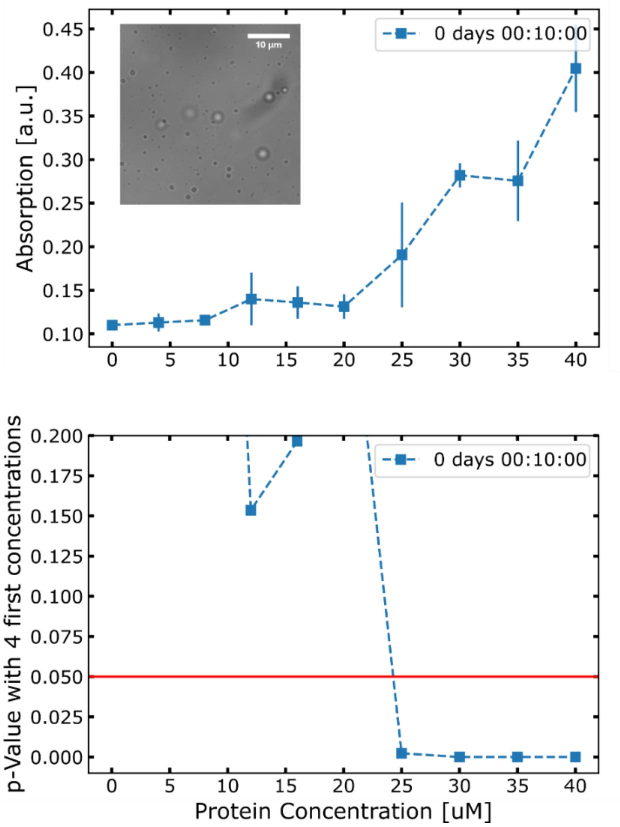


Fig. S4: LLPS are formed at 3M NaCl and probed by absorption. (upper panel) raw absorption data for tau187-WT at different concentration. (lower panel) T-test used to defined the saturation concentration for LLPS formation.

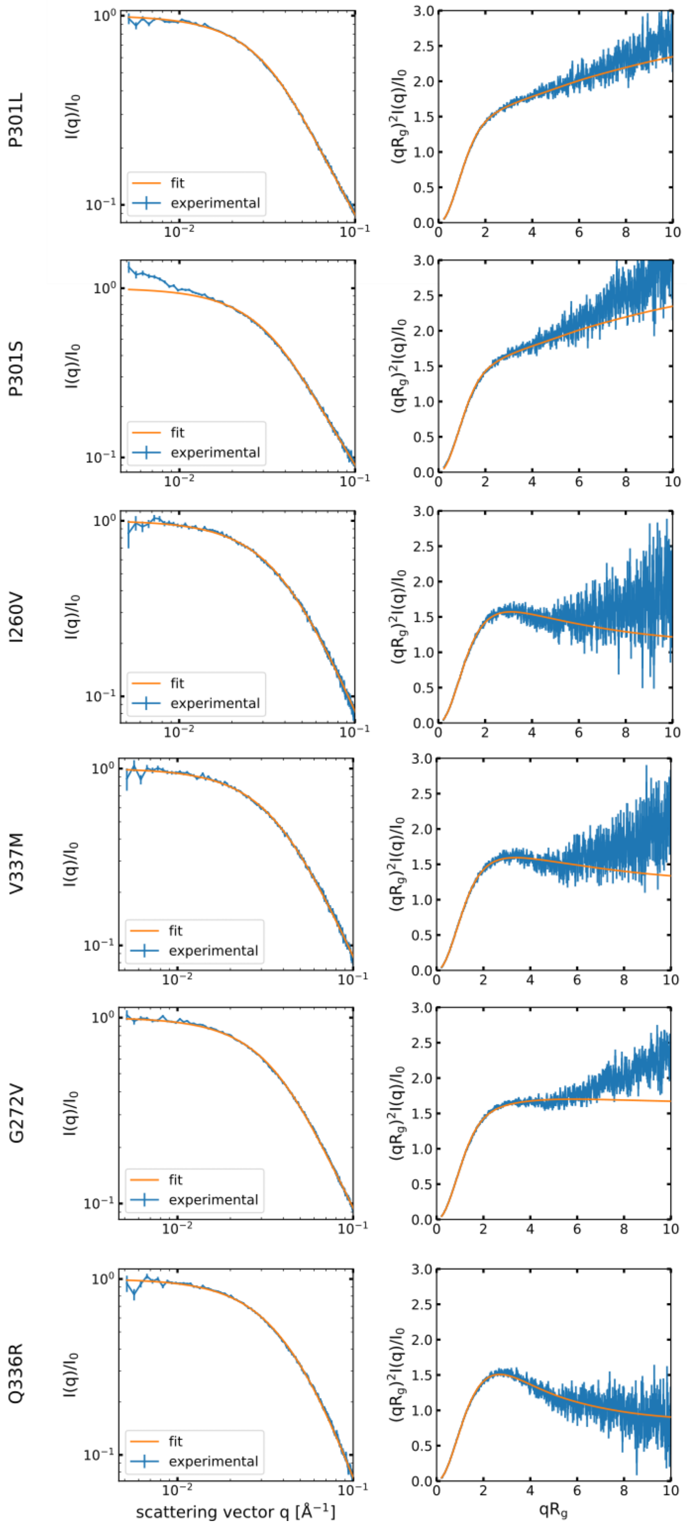


Figure S5 : Best fit of the SAXS data obtained from the server http://sosnick.uchicago.edu/SAXSonIDPs.

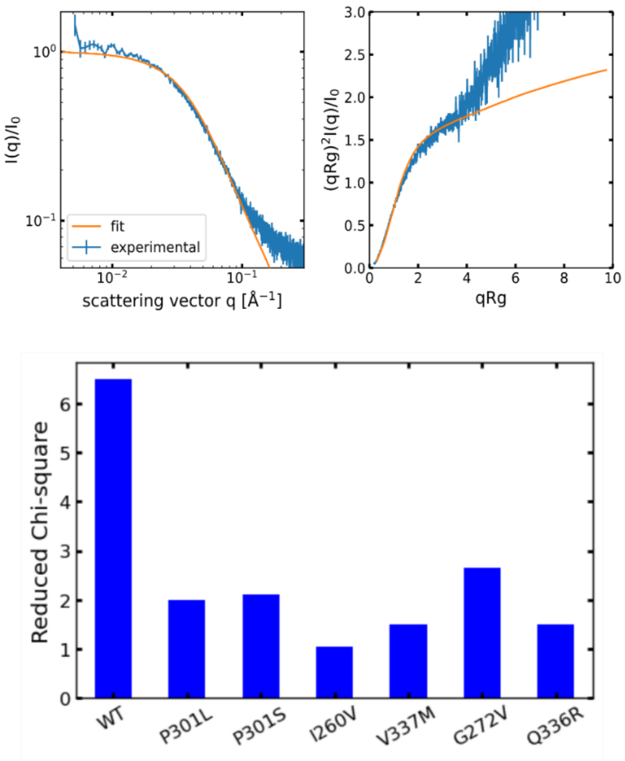


Figure S6 : (Upper panel) Best fit of tau187-WT data from the server http://sosnick.uchicago.edu/SAXSonIDPs. (Lower panel) Reduced chi-square for the fit of the SAXS of each tau mutant. Because the fit was poor for tau187-WT, the output parameters (Flory exponent) were not analyzed in manuscript.

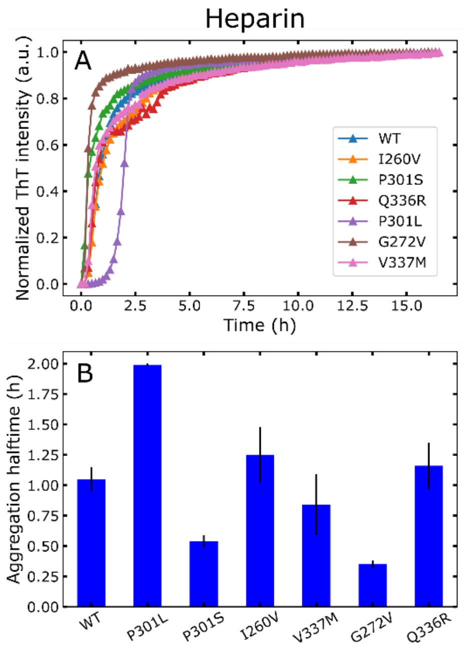


Figure S7 : Heparin suppresses aggregation lag time for almost all mutants, including tau187-WT (A). Aggregation halftime (B) does not correlate with Rg.

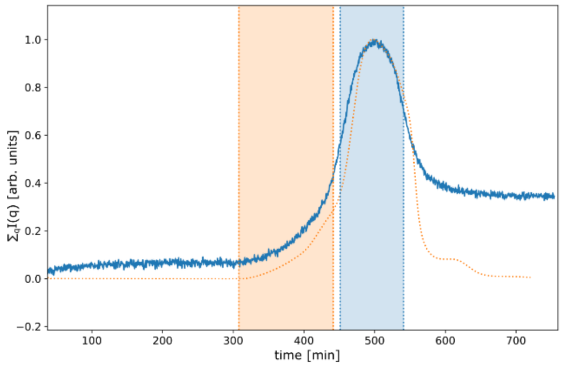


Figure S8: Typical selected regions for the SEC-SAXS data. The total scattering signal is plotted with a solid blue line, the UV measurement is plotted with a dotted orange line with a time delay such that it takes into account the time for the sample to flow from the column output to the X-ray capillary. The selected frames that are averaged for the sample are represented by the blue area and the frames used for the buffer by the orange area.
